# A LRRK2/dLRRK-mediated lysosomal pathway that contributes to glial cell death and DA neuron survival

**DOI:** 10.1101/2021.11.25.469972

**Authors:** Linfang Wang, Honglei Wang, Margaret S. Ho

## Abstract

Mutations in *leucine-rich repeat kinase 2* (*LRRK2*) are the most common cause of familial and sporadic Parkinson’s disease (PD). A plethora of evidence has indicated a role for LRRK2 in endolysosomal trafficking in neurons, while LRRK2 function in glia, although highly expressed, remains largely unknown. Here we present evidence that LRRK2/dLRRK mediates a glial lysosomal pathway that contributes to the mechanism of PD. Independent of its kinase activity, glial LRRK2/dLRRK knockdown in the immortalized microglial cells or flies results in enlarged and swelling lysosomes fewer in number. These lysosomes are less mobile, wrongly acidified, and exhibit defective membrane permeability and reduced activity of the lysosome hydrolase cathespin B. In addition, microglial LRRK2 depletion causes increased *Caspase* 3 levels, leading to glial apoptosis, dopaminergic neurodegeneration, and locomotor deficits in an age-dependent manner. Taken together, these findings demonstrate a functional role of LRRK2/dLRRK in regulating the glial lysosomal pathway; deficits in lysosomal biogenesis and function linking to glial apoptosis potentially underlie the mechanism of DA neurodegeneration, contributing to the progression of PD.

## Introduction

Parkinson’s disease (PD) is the second most common neurodegenerative disorder that exhibits a progressive loss of dopaminergic (DA) neurons and accumulation of Lewy body (LB) protein aggregates in the brains. Given these pathological hallmarks, PD patients often suffer from locomotor dysfunction such as resting tremor, muscle rigidity, and bradykinesia, along with a number of non-motor and cognitive symptoms (Cuenca et al., 2018; Hayes, 2019; Sveinbjornsdottir, 2016). Accumulating evidence indicates that mutations in *leucine-rich repeat kinase 2* (*LRRK2*) are the most common cause of familial and sporadic PD (Di Fonzo et al., 2005; Paisan-Ruiz et al., 2004; Zimprich et al., 2004). LRRK2 is a large multidomain protein (2527 amino acids) that contains the N-terminal domains of armadillo repeat, ankyrin repeat and leucine-rich repeat, and the center enzymatic domains of a Ras-of-Complex (ROC) GTPase, a C-terminal-of-Roc (COR) and a kinase domain, followed by WD40 repeats at its C-terminus. Most LRRK2-associated pathogenic PD mutations including R1441C/G/H, Y1699C, and G2019S are located in the catalytic core (Aasly et al., 2010; Cogo et al., 2020; Erb and Moore, 2020; Haugarvoll et al., 2008; Lin et al., 2008; Rudenko and Cookson, 2014), causing an increase and/or a decrease in the LRRK2 kinase and/or GTPase activity, respectively (Greggio et al., 2006; West et al., 2005; West et al., 2007). Increased LRRK2 kinase activity enhances the phosphorylation of its downstream substrates including a subset of Rab proteins, small GTPases that localize on specific intracellular compartments in the endolysosomal pathway. It is now widely recognized that enhanced LRRK2 kinase activity and its effect on the interacting Rab GTPases contribute to the cause and subsequent progression of PD (Sheng et al., 2012; Steger et al., 2016).

A plethora of evidence reveals increased LRRK2 kinase activity and endolysosomal pathology in DA neurons in the substantia nigra (SN) brain region of PD patients. These pathological alternations include the accumulation of early endosomes and the depletion of late endosomes and lysosomes (Di Maio et al., 2018; Rocha et al., 2020). Lack of LRRK2 in tissues like kidneys expressing LRRK2 in higher levels leads to the formation of large vacuoles and protein aggregates (Herzig et al., 2011; Tong et al., 2012; Tong et al., 2010), increased levels of lysosomal proteins LAMP1, LAMP2 and cathepsin B (cathB) (Baptista et al., 2013; Beilina et al., 2020; Kuwahara et al., 2016; Tong et al., 2012), and perturbation in autophagy (Hinkle et al., 2012; Tong et al., 2010). Furthermore, double knockout mice of *LRRK1* and *LRRK2* exhibit shortened lifespan, as well as mild dopaminergic (DAergic) neuron degeneration and formation of large autophagic vacuoles in the brains (Giaime et al., 2017). These findings implicate a pivotal role for LRRK2 in the endolysosomal pathway pertaining to PD. Of note, LRRK2 kinase activity is also involved in regulating the lysosome number and morphology in primary astrocytes, a type of major brain cells that support neuron growth and participate in various disease pathology (Henry et al., 2015). It has become increasingly clear that LRRK2 activity and the endolysosomal pathology are closely linked, yet whether a direct causal relationship exists, particularly in the brain cell glia, remains largely unexplored.

Given that enhanced LRRK2 kinase activity is a possible cause of PD, targets related to optimized LRRK2 kinase activity or reduced *LRRK2* expression have been proposed as potential therapeutical solutions. It is therefore important to understand the consequence of reducing *LRRK2* expression in physiological and pathological contexts. Here we present evidence that LRRK2 or its *Drosophila* homolog dLRRK regulates the glial lysosomal pathway. Lack of glial *LRRK2/dLRRK* expression causes enlarged and swelling lysosomes in both flies and the immortalized microglial (IMG) cells. These lysosomes devoid of LRRK2/dLRRK are not properly acidified and show signs of defective membrane permeability and disrupted cathB activity, indicating possible leakage of lysosomal content to the cytosol. Interestingly, reducing glial *dLRRK* expression causes locomotor dysfunction and DA neurodegeneration in an age-dependent manner, both parkinsonian symptoms widely recognized by a *Drosophila* PD model. Taken together, our findings illustrate a role for LRRK2/dLRRK in the glial lysosomal pathway pertaining to PD.

## Results

### Reducing *dLRRK* expression in glia causes enlarged lysosomes fewer in number

To investigate the consequence of reducing *LRRK2/dLRRK* expression in glia, the transgenic RNAi line targeting *dLRRK* was expressed using a pan-glial driver, *repo-Gal4. dLRRK* expression was efficiently silenced by RNAi as assessed by qRT-PCR (Figure S1A). Using Lamp1.GFP as a marker, lysosomes were labeled as GFP-positive puncta in membranous RFP-marked glial cells (*repo*>*UAS-mCD8.RFP*, *UAS-Lamp1.GFP*). Regions analyzed in the adult brains are enriched of glial nuclei, neuronal nuclei, and glial processes as illustrated in grey in the schematic diagram (Figure 1A). Interestingly, Lamp1.GFP-labeled glial lysosomes were enlarged and exhibited swelling morphology when expressing *dLRRK* RNAi (second panel, Figure 1D). These lysosomes exhibited an increase in the size and a decrease in the number (Figures 1B and 1C), and are less mobile (Figures S1C and S1D). The enlarged lysosomes were progressively bigger in adult flies from 3-to 20-day-old, suggesting that the lysosomes might deteriorate in an age-dependent manner. *dLRRK* overexpression did not cause significant change in glial lysosome structure and morphology in young and old adult flies, whereas reintroducing *dLRRK* in adult fly glia expressing *dLRRK*-RNAi rescued the lysosomal defects (third and fourth panels, Figure 1D). It is noteworthy to mention that Rab7.GFP-positive late endosomes lacking glial dLRRK were also enlarged (Figures S1E and S1F) and less mobile (Figures S1G and S1H), indicating a general endolysosomal defect when glial dLRRK is absent.

**Figure 1.**
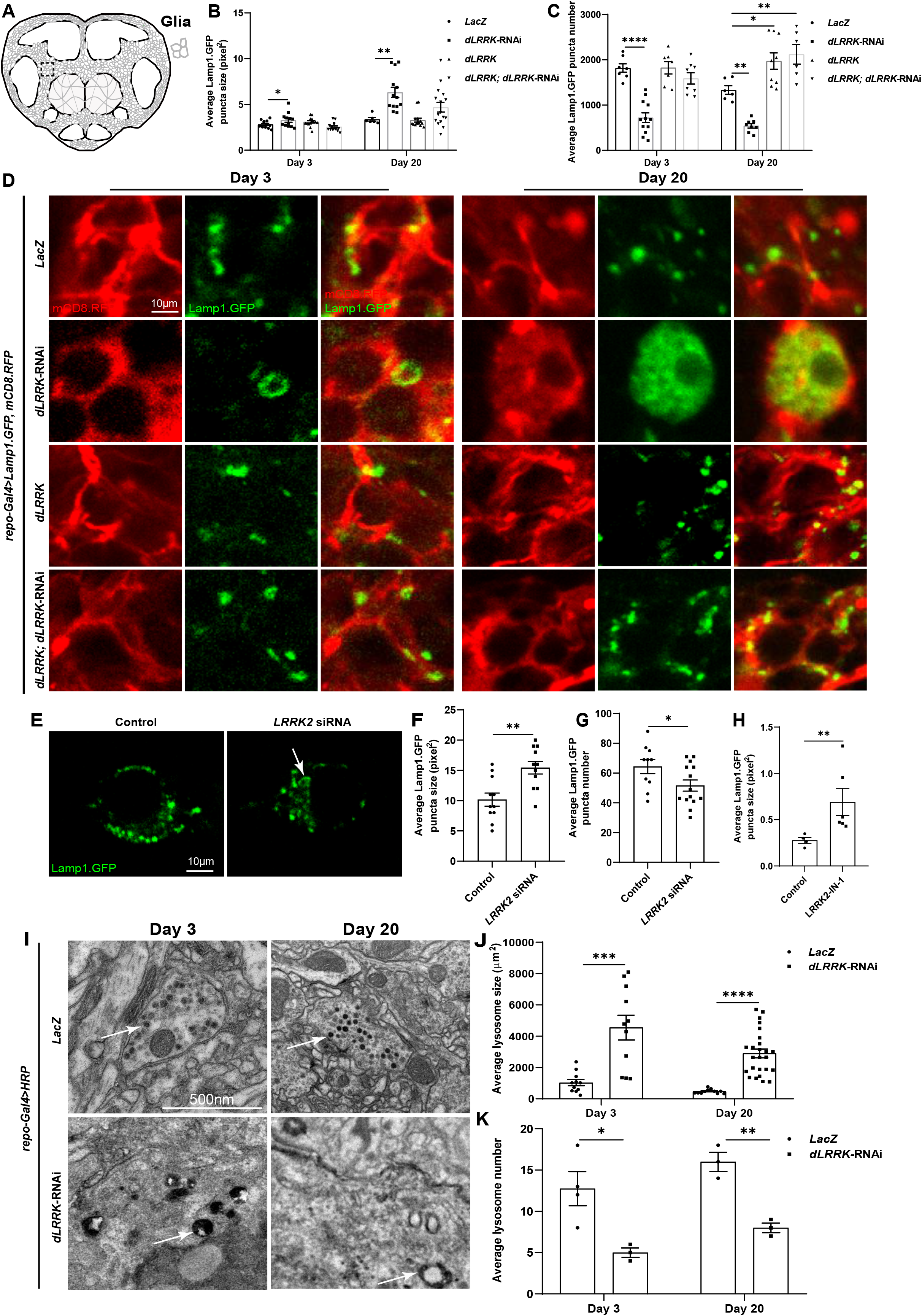
Lack of glial LRRK2/dLRRK causes enlarged lysosomes fewer in number. (**A**) Schematic diagram of an adult fly brain. Areas in grey are where glial nuclei, neuronal nuclei, and glial processes reside, whereas areas in white represent major axon neuropils with glial wrapping. Black square indicates where images were captured in D. (**B** and **C**) Quantifications of D. Average lysosome size and number represented by the Lamp1.GFP puncta are quantified. (**D**) Representative images of glial lysosomes in adult fly brains with altered *dLRRK* expression. Glial lysosomes (green) are labeled with *repo-GAL4* driven *UAS-Lamp1.GFP* while glial processes are marked with *UAS-mCD8.RFP* (*repo-Gal4*>*UAS-Lamp1.GFP*, *UAS-mCD8.RFP*). Note that downregulating glial *dLRRK* induces the formation of enlarged lysosomes with decreased number. Genotypes analyzed include: *repo-Gal4* driven *UAS-LacZ*, *UAS-dLRRK*-RNAi, *UAS-dLRRK*, and *UAS-dLRRK*; *UAS-dLRRK*-RNAi. (**E-H**) Representative images (E) and quantifications (F-H) of microglial lysosomes in IMG cells treated with the *LRRK2* siRNA or inhibitor IN-1. Note that these lysosomes devoid of LRRK2 are also enlarged and reduced in number. (**I**-**K**) Representative TEM images (I) and quantifications (J and K) of lysosomes in the control and *repo*>*dLRRK*-RNAi adult fly brains. Note an increase in the size and a decrease in the number of lysosomes (white arrows in I) when *dLRRK*-RNAi is expressed in glia. Scale bars: 10 μm for D, E and 500 nm for I. For adult fly brains or IMG cells, quantifications are done by averaging the intensities, size, or number per acquired brain region or per cell, respectively. For TEM images, average 4 independent fly brains of each genotype were analyzed. Quantifications are done by averaging the lysosome number per brain region. For lysosome size in the TEM image, average lysosome size was quantified by measuring the size of all lysosomes in all brain regions. Size per lysosome was shown on the scatter plot (F). Sample number n is indicated by the dot number in the scatter plot. Data are shown as mean ± SEM. P-values of significance (indicated with asterisks, ns no significance, * p<0.05, ** p<0.01, and *** p<0.001) are calculated by two-tailed unpaired t-test or ordinary one-way ANOVA followed by Tukey’s multiple comparisons test. Approximately 15 (adult fly brains) or 10 (IMG cells) confocal Z-stack sections were taken with 0.6-1 μm each, and representative single layer images acquired at the similar plane of brains across all genotypes are shown. Pearson’s R values for colocalization are calculated based on methods described. All data in subsequent Figures are present and analyzed as described in Figure 1 unless stated otherwise.

To demonstrate that dLRRK function in the glial lysosomal pathway is conserved in vertebrates, we turned to the mouse immortalized microglial (IMG) cells (McCarthy et al., 2016). Following siRNA treatment, which efficiently silenced *LRRK2* expression (Figure S1B), IMG cells were transfected with the Lamp1.GFP plasmid to label the lysosomes. Consistent with the adult fly brains, lysosomes of IMG cells treated with the *LRRK2* siRNA or inhibitor LRRK2-IN-1 (Deng et al., 2011; Munoz et al., 2015) were enlarged and fewer in number (Figures 1E–1H). Furthermore, Transmission Electron Microscopy (TEM) analysis also revealed that lysosomes were enlarged with abnormal morphology and fewer in number upon glial dLRRK depletion in adult flies of 3- and 20-day-old (arrows, Figures 1I–1K). Taken together, these results suggest that LRRK2/dLRRK regulates glial lysosomal biogenesis.

### Enlarged lysosomes devoid of LRRK2/dLRRK are wrongly acidified

To further investigate the properties of these enlarged lysosomes, Lysotracker Red (LTR) dye was used to detect the lysosome acidity *in-vivo*. Interestingly, the mean LTR intensity of lysosomes co-positive for Lamp1 and LTR decreased significantly upon glial dLRRK depletion in adult fly brains of 3- and 20-day-old (Figures 2A and 2B). Lamp1-LTR colocalization value was also evidently lowered, suggesting that these enlarged lysosomes are not as acidic (Figure 2C). Similarly, Lamp1-positive microglial lysosomes treated with *LRRK2* siRNA or inhibitor exhibited reduction in the mean LTR intensities and Lamp 1-LTR colocalization (Figures 2D–2H). Taken together, these results indicate that lack of LRRK2/dLRRK in glia triggers the formation of enlarged lysosomes that are wrongly acidified; these swelling lysosomes with abnormal acidity are likely dysfunctional.

**Figure 2.**
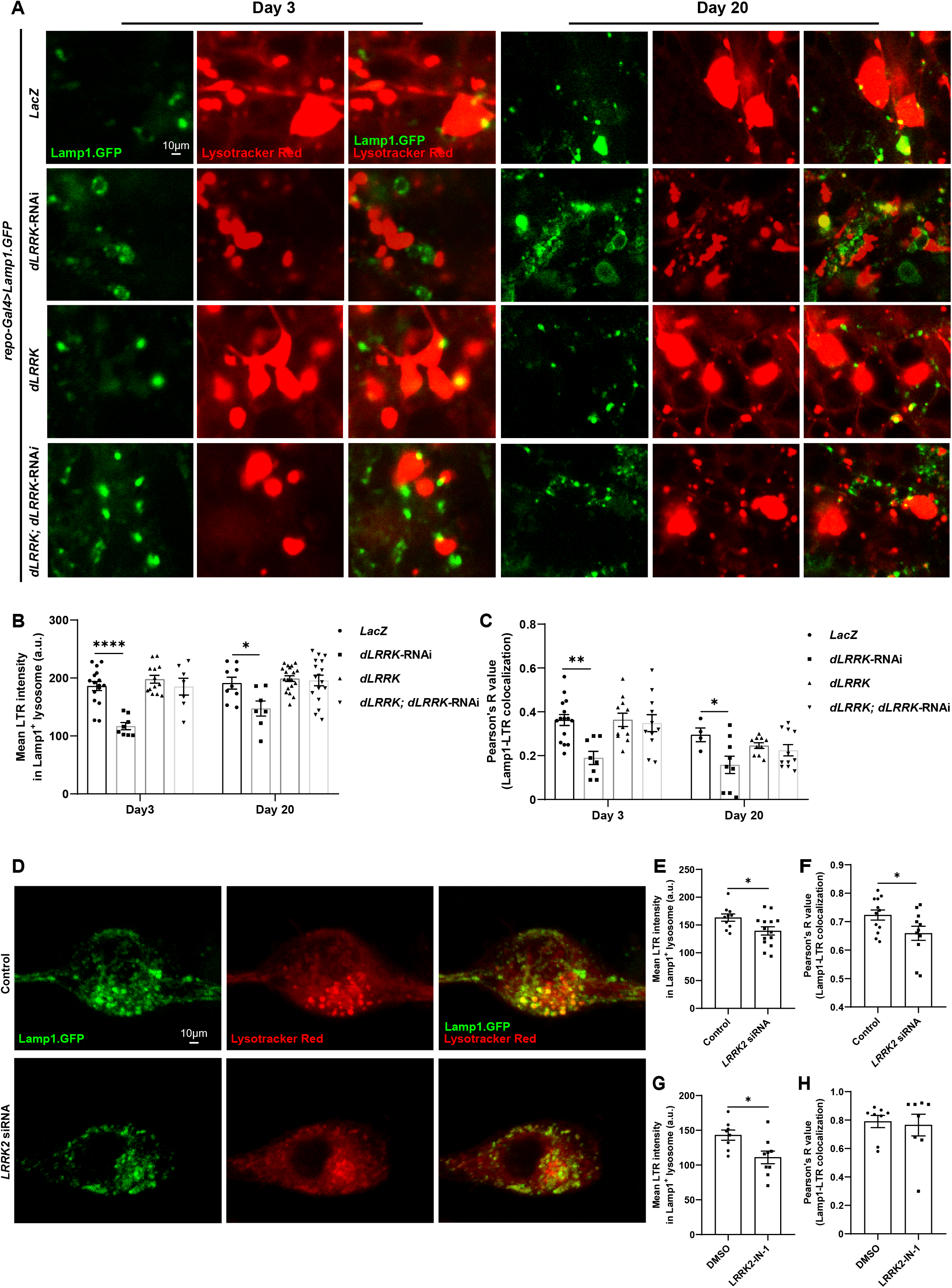
Glial lysosomes devoid of LRRK2/dLRRK are dysfunctional. (**A-C**) Representative images (A) and quantifications (B and C) of glial lysosomes staining with Lysotracker Red (LTR) dye in adult fly brains with altered *dLRRK* expression. Glial lysosomes (green) are labeled with *repo-GAL4* driven *UAS-Lamp1.GFP* (*repo-Gal4*>*UAS-Lamp1.GFP*). The lysosome acidity is detected with LTR dye in red. Note that downregulating glial *dLRRK* induces the formation of enlarged lysosomes, which are abnormally acidic. Genotypes analyzed include: *repo-Gal4* driven *UAS-LacZ*, *UAS-dLRRK-RNAi*, *UAS-dLRRK*, and *UAS-dLRRK*; *UAS-dLRRK*-RNAi. (**D**-**H**) Representative images (D) and quantifications (E-H) of microglial lysosomes in IMG cells treated with the *LRRK2* siRNA (D-F) or inhibitor (G and H). Note that these lysosomes devoid of LRRK2 are wrongly acidified. Scale bars and the quantified sample number n are indicated as in Figure 1. Data are shown as mean ± SEM. P-values of significance (indicated with asterisks, ns no significance, * p<0.05, ** p<0.01, and *** p<0.001) are calculated by two-tailed unpaired t-test or ordinary one-way ANOVA followed by Tukey’s multiple comparisons test.

### Microglial LRRK2 mediates lysosome membrane permeabilization

Given that lysosome swelling precedes and is indicative of defects in lysosome membrane permeabilization (LMP) (Hasan et al., 2008; Wang et al., 2018), we tested if microglial LRRK2 regulates lysosome membrane permeability. Interestingly, Acrid Orange (AO) staining revealed that red and green fluorescent intensity decreased and increased, respectively upon microglial LRRK2 depletion by siRNA or inhibitor, indicating the lysosomal pH gradient and membrane permeability are disrupted (Figures 3A–3D). Treatment with the Lysotracker Green (LTG) dye also revealed a clearly diffuse pattern with hollow puncta, suggesting possible leakage of lysosomal content into the cytosol (Figure 3E). Quantifications revealed that these hollow puncta are bigger in size (Figure 3F) and exhibited lower and higher LTG fluorescent signals inside and outside of the puncta, respectively (Figures 3G). Results from a parallel experiment co-transfecting the Lamp1-RFP as a lysosome marker also indicated that the Lamp1-positive puncta are bigger in size (Figure 3H) and exhibited lower LTG intensity (Figure 3I). In addition, Magic Red staining revealed a reduction in the cathB activity inside the Lamp1-positive lysosomes and the cathB-Lamp1 colocalization in IMG cells treated with the *LRRK2* siRNA or inhibitor (Figures 3J–3N). These results demonstrate that microglial lysosomes devoid of LRRK2 exhibit defective membrane permeability.

**Figure 3.**
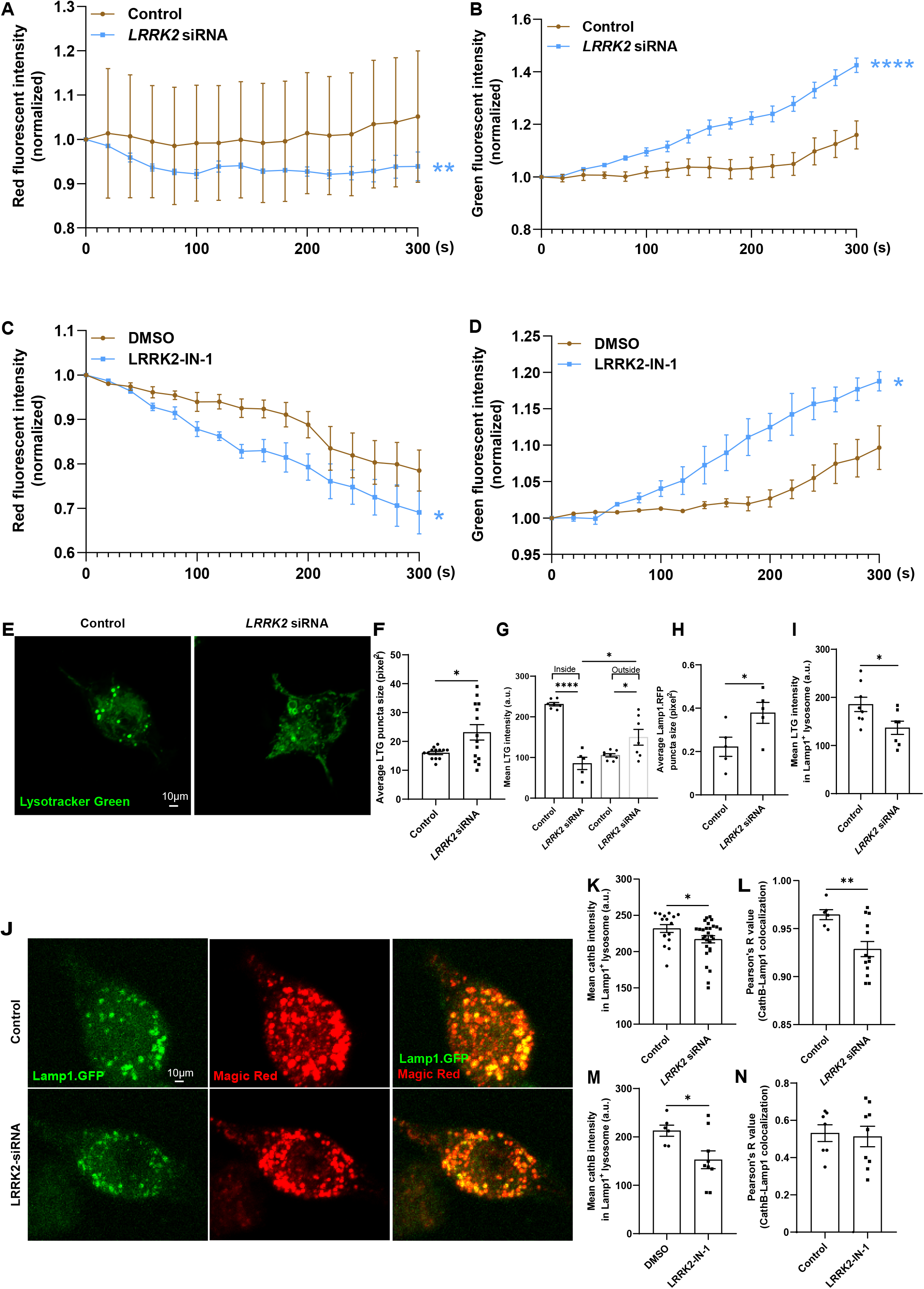
Microglial lysosomes devoid of LRRK2 exhibit defective membrane permeabilization. (**A-D**) Quantification curves of AO staining on IMG cells treated with the *LRRK2* siRNA or inhibitor. Note that the red and green fluorescent intensity are significantly lowered and higher, respectively over the time course analyzed upon microglial LRRK2 depletion. Average 10 cells were quantified for each experiment. (**E-H**) Representative images (E) and quantifications (F-I) of Lysotracker Green (LTG)-positive puncta in IMG cells treated with the *LRRK2* siRNA. Note that these puncta devoid of LRRK2 are hollow and enlarged. LTG fluorescent signals increase and exhibit a diffuse pattern outside of puncta (with or without Lamp1-RFP marking the lysosomes). (**J-N**) Representative images (J) and quantifications (K-N) of Magic Red staining on IMG cells treated with the *LRRK2* siRNA. Note that Magic Red staining representing cathespin B activity and its colocalization with Lamp1 decreased upon microglial LRRK2 depletion. Scale bars and the quantified sample number n are indicated as in Figure 1. Data are shown as mean ± SEM. P-values of significance (indicated with asterisks, ns no significance, * p<0.05, ** p<0.01, and *** p<0.001) are calculated by two-tailed unpaired t-test or ordinary one-way ANOVA followed by Tukey’s multiple comparisons test.

### Lack of LRRK2/dLRRK increases Caspase 3 expression and glial apoptosis

Given that lysosome swelling and defective LMP frequently connects to cell death, we next tested if glial cell death is affected by the absence of LRRK2/dLRRK. Whereas the expression of components in the inflammasome-mediated pyroptosis pathway including *NOD-like receptor protein 3* (*NLRP3*), *gasdermin D* (*GSDMD*) and *Caspase 1* were all decreased upon microglial LRRK2 depletion, *Caspase 3* expression increased significantly, suggesting that apoptosis is triggered (Figure 4A). Annexin-V staining of IMG cells also indicated an induction of early apoptosis when lacking microglial LRRK2 (Figures 4B and 4C). Moreover, TUNEL staining of IMG cells and adult fly brains revealed elevated number of apoptotic cells when lacking glial LRRK2/dLRRK (Figures 4D–4H). *In toto*, these results suggest that lack of LRRK2/dLRRK induces glial apoptosis, potentially a consequence of the underlying LMP defects.

**Figure 4.**
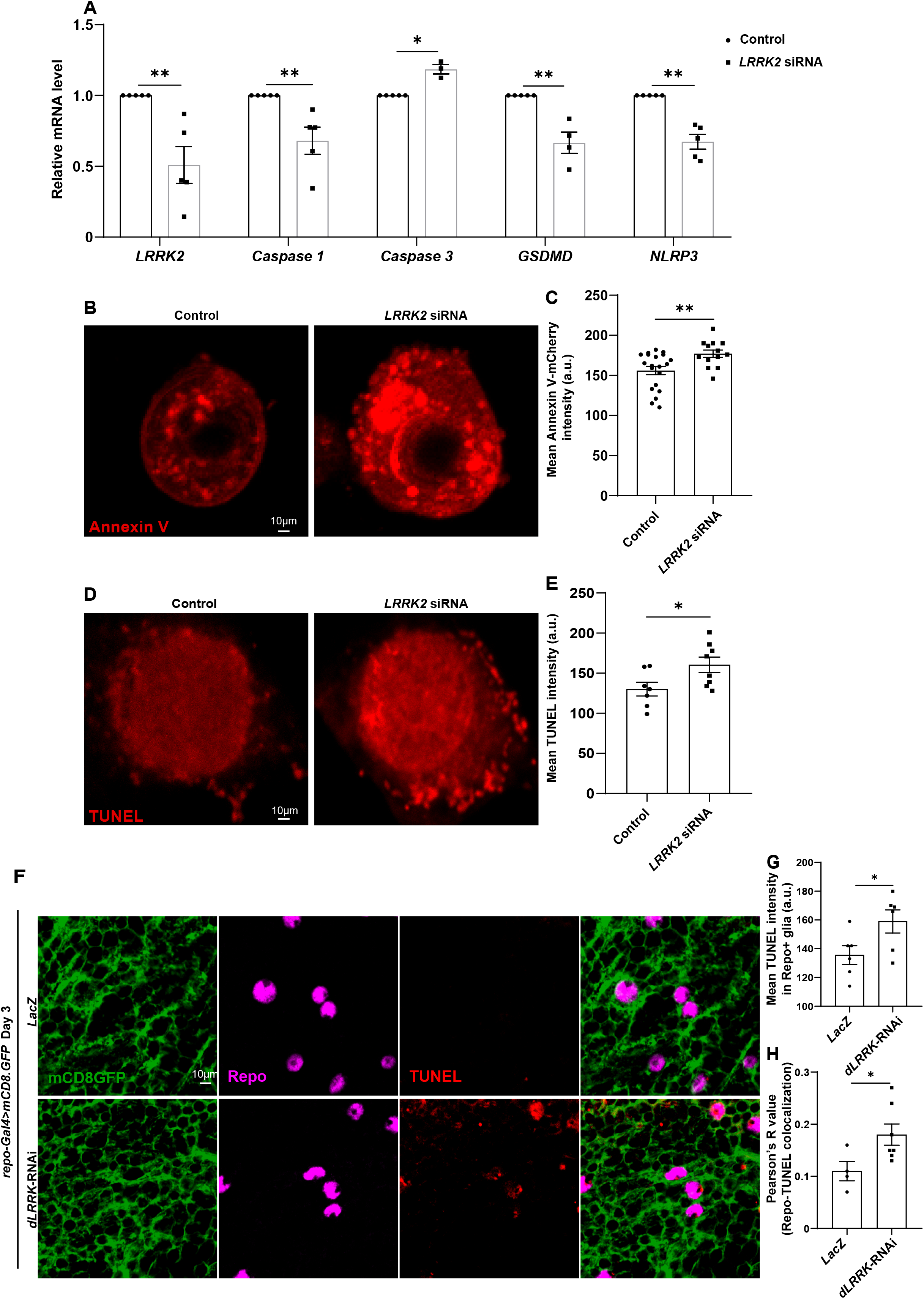
Lack of LRRK2/dLRRK causes increased *Caspase 3* expression and glial apoptosis. (**A**) Relative mRNA levels of genes including *LRRK2*, *Caspase 1*, *Caspase 3*, *GSDMD*, and *NLRP3* were assessed by qRT-PCR upon microglial LRRK2 depletion. Note that only the *Caspase 3* level increases. (**B** and **C**) Representative images (B) and quantifications (C) of IMG cells stained with Annexin-V to detect early apoptotic events. (**D**-**H**) Representative images (D and F) and quantifications (E, G, H) of IMG cells and adult fly brains stained with TUNEL to detect apoptosis. Note that TUNEL staining of apoptotic cells increases upon glial LRRK2/dLRRK depletion. Scale bars and the quantified sample number n are indicated as in Figure 1. Data are shown as mean ± SEM. P-values of significance (indicated with asterisks, ns no significance, * p<0.05, ** p<0.01, and *** p<0.001) are calculated by two-tailed unpaired t-test or ordinary one-way ANOVA followed by Tukey’s multiple comparisons test.

### Glial dLRRK contributes to DA neurodegeneration and locomotor deficits

To further demonstrate the role of glial dLRRK in pathological contexts, DA neurodegeneration and locomotor behavior were analyzed in flies with altered *dLRRK* expression. Fly DA neurons are located in clusters and named upon their relative positions in adult brains (Budnik and White, 1988; Nassel and Elekes, 1992). These clusters include protocerebral posterior lateral (PPL)1, PPL2, protocerebral posterior medial (PPM)1, PPM2, and PPM3 (Figure 5A). Given that the number of PPM1/2 DA neurons has been shown to degenerate upon α-Synuclein overexpression in flies (Feany and Bender, 2000), we quantified the DA neuron number in this cluster when altering *dLRRK* expression in glia. Interestingly, whereas no significant alternation in the number of PPM1/2 DA neurons was detected when overexpressing glial *dLRRK*, these DA neurons deteriorated and exhibited a decrease in number progressively when glial *dLRRK* was absent (Figures 5B and 5C). Co-expressing *dLRRK* in the presence of *dLRRK*-RNAi rescued the decrease in the DA neuron number. These results indicate that glial dLRRK contributes to DA neuron survival. Furthermore, while *dLRRK* overexpression causes a slight increase in the climbing distance, *dLRRK*-RNAi expressing flies exhibited locomotor deficits as shown by the decrease in the climbing distance in an age-dependent manner (Figures 5D and 5E). These results indicate that glial dLRRK regulates fly locomotor activity, further supporting the notion that glial dLRRK contributes to PD-like symptoms in flies.

**Figure 5.**
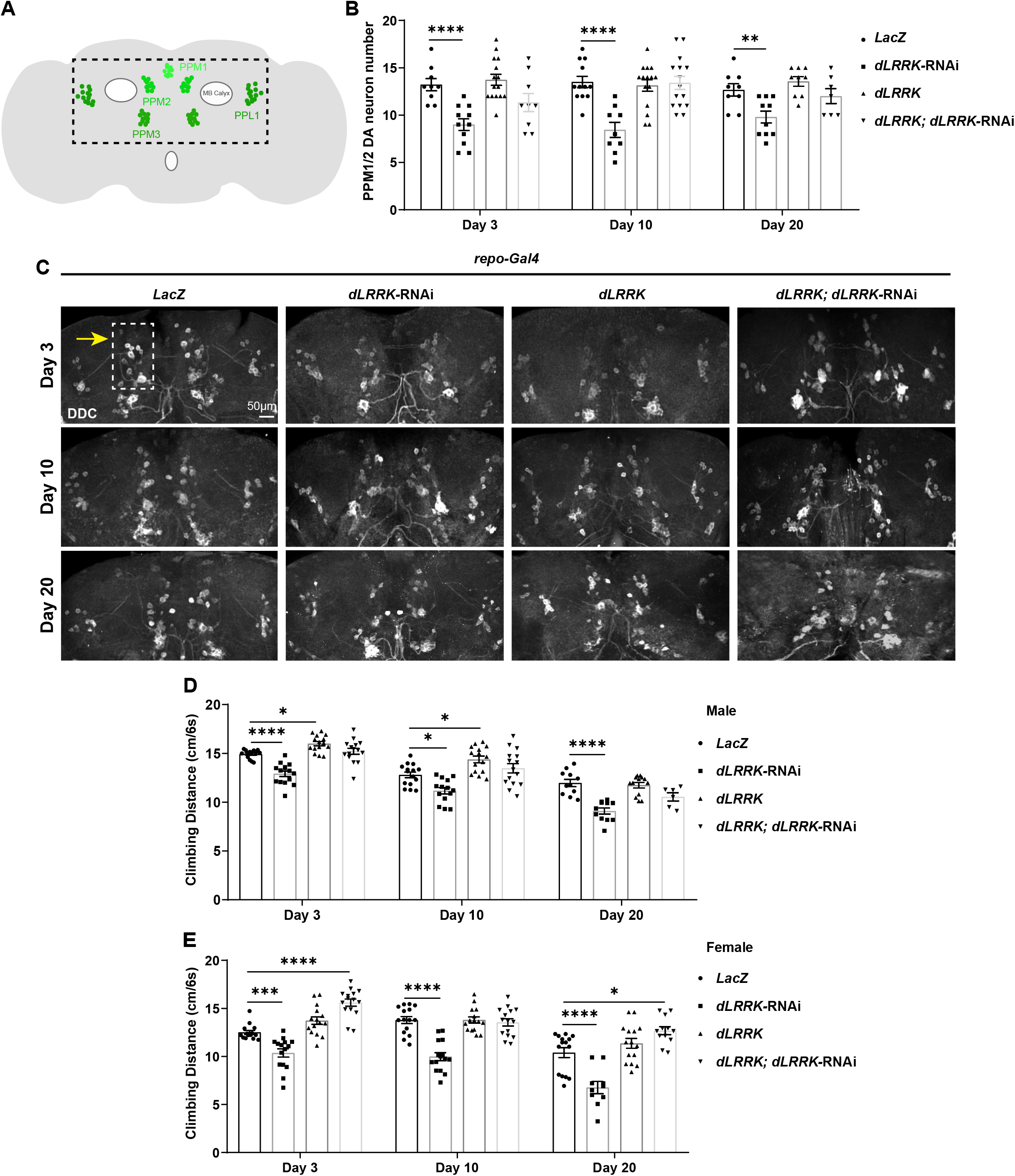
Lack of glial dLRRK causes DA neurodegeneration and locomoter deficits. (**A**) Illustration of an adult fly brain with different DA neuron clusters (green). Note that for simplicity only PPM1/2, PPM3, and PPL1 clusters are shown. DA neurons in the PPM1/2 cluster are visualized in C and quantified in B. (**B** and **C**) Representative images (B) and quantifications (C) of DA neuron number in the adult fly brains with the indicated genotypes. Note that DDC-positive DA neuron number in the PPM1/2 cluster decreases in an age-dependent manner upon expressing *dLRRK*-RNAi in glia. (**D** and **E**) Analysis of fly locomotor behavior in both males and females reveals a significant decline in the locomotor activity upon glial dLRRK depletion. Genotypes analyzed include: *repo-Gal4* driven *UAS-LacZ*, *UAS-dLRRK*-RNAi, *UAS-dLRRK*, and *UAS-dLRRK*; *UAS-dLRRK-RNAi*. Scale bars and the quantified sample number n are indicated as in Figure 1. Data are shown as mean ± SEM. P-values of significance (indicated with asterisks, ns no significance, * p<0.05, ** p<0.01, and *** p<0.001) are calculated by two-tailed unpaired t-test or ordinary one-way ANOVA followed by Tukey’s multiple comparisons test.

### Enhanced LRRK2 kinase activity does not affect glial lysosome morphology

To further elucidate the loss-of-function mechanism of LRRK2/dLRRK, we next tested whether enhanced LRRK2 kinase activity, the prevalent cause of PD, leads to similar lysosomal or behavioral defects. Unlike *dLRRK*-RNAi, glial expression of human *LRRK2* (*hLRRK2*) caused a reduction in the lysosome size in adult flies of 3- and 20-day-old (Figures 6A and 6B). The glial lysosome number also increased in *hLRRK2-expressing* adult flies of 20-day-old (Figure 6C). These results are opposite to the ones observed when downregulating *dLRRK2* expression in glia, suggesting that glial *dLRRK* expression needs to be precisely regulated. Moreover, enhanced LRRK2 kinase activity by overexpressing the LRRK2^G2019S^ variant in flies did not cause severe alternation in lysosome size nor number (Figures 6A–6C), suggesting that dLRRK mediates the glial lysosomal pathway independent of its kinase activity. Despite so, overexpression of *hLRRK2* or *LRRK2^G2019S^* induces DA neurodegeneration (Figures 6D and 6E), validating our animal PD model and further demonstrating that glial dLRRK contributes to the PD-like symptoms in flies.

**Figure 6.**
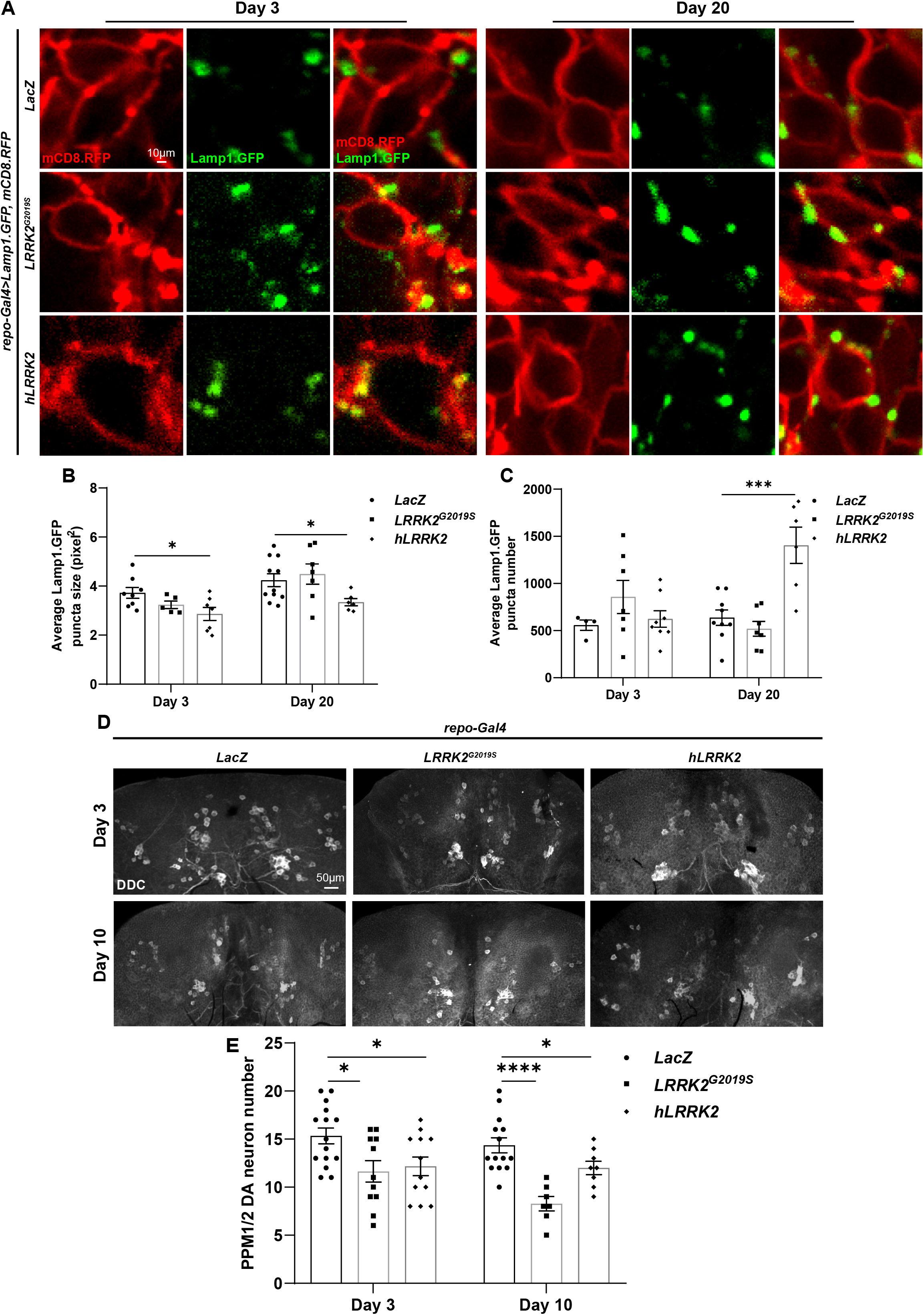
Enhanced LRRK2 kinase activity causes DA neurodegeneration independent of lysosomal homeostasis. (**A-C**) Representative images (A) and quantifications (B and C) of glial lysosomes in adult fly brains with exogenous *LRRK2* expression. Glial lysosomes (green) are labeled with *repo-GAL4* driven *UAS-Lamp1.GFP* while glial processes are marked with *UAS-mCD8.RFP* (*repo-Gal4*>*UAS-Lamp1.GFP*, *UAS-mCD8.RFP*). Note that overexpressing glial *hLRRK2* causes a reduction in the lysosome size (B) and an increase in the lysosome number (C), opposite to what have been observed when expressing *dLRRK*-RNAi. *LRRK2^G2019S^* overexpression does no induce the formation of enlarged lysosomes. Genotypes analyzed include: *repo-Gal4* driven *UAS-LacZ*, *UAS-LRRK2^G2019S^*, and *UAS-hLRRK2*. (**D** and **E**) Representative images (D) and quantifications (E) of DDC-positive DA neuron number in the adult fly brains with the indicated genotypes. Note that DA neuron number in the PPM1/2 cluster decreases upon expressing *LRRK2^G2019S^* or *hLRRK2* in glia. Scale bars and the quantified sample number n are indicated as in Figure 1. Data are shown as mean ± SEM. P-values of significance (indicated with asterisks, ns no significance, * p<0.05, ** p<0.01, and *** p<0.001) are calculated by two-tailed unpaired t-test or ordinary one-way ANOVA followed by Tukey’s multiple comparisons test.

## Discussion

Accumulating evidence has now indicated that enhanced LRRK2 kinase activity is a possible cause of PD. Although concomitantly seen in the brains of PD patients, whether the enhanced LRRK2 kinase activity is a direct cause of endolyososomal pathology remains unknown. Our findings indicate that these two events might be independent from each other, as enhanced LRRK2 kinase activity does not cause lysosome abnormality in *Drosophila* glia. Intriguingly, consistent to previous observations, reduced *LRRK2/dLRRK* expression causes enlarged and swelling lysosomes, indicating that LRRK2/dLRRK loss-of-function might be a prominent factor in generating endolysosomal pathology in PD; our findings provide a different perspective in understanding the mechanism of LRRK2/dLRRK causing PD.

Despite that glia are important modulators of PD progression, LRRK2/dLRRK function in glia have not been largely explored. LRRK2 is highly expressed in mouse astrocytes and human microglia (Zhang et al., 2014; Zhang et al., 2016) and enhanced LRRK2 kinase activity has been shown to regulate lysosome number and morphology in primary astrocytes (Henry et al., 2015). Our findings further implicate a role for LRRK2 in microglia. LRRK2 regulates the number and size of microglial lysosomes, lysosome membrane permeability, and microglial apoptosis. These results echo observations in *Drosophila* glia, both supporting the notion that LRRK2/dLRRK plays a pivotal role in the glial lysosomal pathway. Considering the importance of glial contribution to PD, it is likely that LRRK2 might regulate disease progression via a glial mechanism involving lysosomal homeostasis.

The enlarged lysosomes are less mobile, abnormally acidic, and exhibit defective membrane permeability, which potentially leads to the leakage of lysosomal content to the cytosol. It has been shown that lysosome swelling precedes LMP, and depends on the degree of LMP disruption, partial or complete cell death occurs (Hasan et al., 2008; Wang et al., 2018). Under partial LMP, the TFEB-dependent survival mechanism is activated, so that both TFEB transcription and lysosome biogenesis are promoted. However, in our case of microglial LRRK2 depletion, TFEB and TFE3 transcription are downregulated (Figure S1I), suggesting inactive reformation of lysosomes and severe cell death. Furthermore, lacking glial LRRK2/dLRRK not only disrupts LMP, provokes glial apoptosis, but also causes DA neurodegeneration, indicating that these glial events are closely linked to neuronal survival. Given that lysosome-related dysregulation and cell death have been detected in cellular and animal models of PD, our findings nicely recapitulate these defects upon glial LRRK2/dLRRK depletion, identifying the LRRK2/dLRRK-mediated lysosomal pathway as the underlying mechanism of glial cell death and DA neurodegeneration.

## Acknowledgments

We thank Bloomington *Drosophila* Stock Center, Vienna *Drosophila* RNAi Center, Tsinghua Fly Center, the Core Facility of *Drosophila* Resource and Technology, Shanghai Institute of Biochemistry and Cell Biology, Chinese Academy of Sciences, Developmental Studies Hybridoma Bank, Cheng-Ting Chien, Yijun Liu, and Aike Guo for fly stocks and antibodies; Chih-Hao Lee for the IMG cell line. We also thank the Molecular Imaging Core Facility (MICF), the Molecular and Cell Biology Core Facility (MCBCF), and the Multi-Omics Core Facility (MOCF) at the School of Life Science and Technology, ShanghaiTech University for providing technical support; Yu Kong and Lijun Pan in Electron Microscopy Facilities of Center for Excellence in Brain Science and Technology, Chinese Academy of Science for assistance with EM sample preparation. We thank Ho lab members for discussion and comments. This work was supported by grants from ShanghaiTech, National Natural Science Foundation of China (31871039 and 32170962), and Shanghai High-end Foreign Expert Program (#21WZ2502300).

## Author contributions

L.W and M.S.H conceived and designed the study. L.W and H.W performed the experiments. L.W and M.S.H analyzed the data. M.S.H wrote the paper with the input of L.W. All authors read and approved the manuscript.

## Materials and Methods

### *Drosophila* genetics

Flies were raised under standard conditions at 25°C with 70% humidity. All fly crosses were carried out at 25°C with standard laboratory conditions unless noted otherwise. All strains were obtained from Bloomington *Drosophila* Stock Center (BDSC), the Vienna *Drosophila* RNAi Center (VDRC), Tsinghua Fly Center, or as gifts from colleagues. The following fly strains were used in this study: isogenic *repo-GAL4* (a pan-glial driver), *UAS-mCD8.GFP*, *UAS-mCD8.RFP*, *UAS-LacZ* (BL1777), *UAS-dLRRK-RNAi* (BL39019), *UAS-dLRRK* (BL34750), *UAS-LRRK2^G2019S^* and *UAS-hLRRK2* (Cheng-Ting Chien), *UAS-Lamp1.GFP*, *UAS-Rab7.GFP* (BL42705), *UAS-HRP* (Aike Guo).

### Cell line and transfection

IMG cell line was a gift from Chih-Hao Lee (McCarthy et al., 2016). Lamp1-GFP and Lamp1-RFP plasmids were gifts from Yijun Liu (Zhejiang University). Cells were cultured in Dulbecco’s modified Eagle medium (Cat. #11965092, Gibco, New York, NY, USA) with high glucose (4.5 g/L), 10 % fetal bovine serum (Cat. #10099141, Gibco) and 100 units/mL penicillin-streptomycin (Cat. #15140-122, Gibco) at 37°C with 5% CO2 humidity. IMG cells were transfected using Lipo2000 (Cat. #11668019, Life Technologies, Carlsbad, CA, USA) according to the manufacturer’s protocol. In brief, cells were seeded on a 12-well plate overnight. 40 pmol siRNA (GenePharma), 500 ng plasmid and 2 ul Lipo 2000 were diluted with 100 μl Opti-MEM (Cat. #31985070, Gibco), mixed, and incubated at room temperature for 20 minutes, then added dropwise to cells cultured at 37°C with 5% CO2. After 6 hours, cells were changed into growth medium and harvested after 72 hours. For LRRK2 inhibition, IMG cells were treated with the LRRK2 catalytic inhibitor LRRK2-IN-1 (Cat. #1234480-84-2, Selleck, Houston, HOU, USA) at 5uM for 2 hours.

### Molecular biology

For qRT-PCR, total RNAs were extracted from adult fly heads or IMG cells using TransZol Up (Cat. #ET111-01, TransGen, Beijing, China). cDNAs were reversely transcribed using HiScript III RT SuperMix (Cat. #R323-01, Vazyme, Nanjing, China). qRT-PCR reactions were performed using ChamQ Universal SYBR qPCR Master Mix (Cat. #Q711-02, Vazyme) and ABI 7500 RT-PCR system. The mRNA levels were normalized to *rp49* (flies) or *Actin* (IMG cells). The expression levels were analyzed by ΔΔCT method. Primers used are listed below:

Flies:

*rp49*-F: CCACCAGTCGGATCGATATGC

*rp49*-R: CTCTTGAGAACGCAGGCGACC

*dLRRK*-F: CCGCTTGTTCCGTTGTTGTG

*dLRRK*-R: ATCTTTCCTGCAATTTCGC

IMG cells:

*Actin*-F: CTAAGGCCAACCGTGAAAAG

*Actin*-R: ACCAGAGGCATACAGGGACA

*LRRK2*-F: ATCTCACCCTTCATGCTTTCTG

*LRRK2*-R: TCTCAGGTCGATTGTCTAAGACT

*Caspase1*-F: ACAAGGCACGGGACCTATG

*Caspase1*-R: TCCCAGTCAGTCCTGGAAATG

*Caspase3*-F: ATGGAGAACAACAAAACCTCAGT

*Caspase3*-R: TTGCTCCCATGTATGGTCTTTAC

*GSDMD*-F: CCATCGGCCTTTGAGAAAGTG *GSDMD*-R: ACACATGAATAACGGGGTTTCC *NLRP3*-F: ATTACCCGCCCGAGAAAGG

*NLRP3*-R: TCGCAGCAAAGATCCACACAG *TFEB*-F: CCACCCCAGCCATCAACAC *TFEB*-R: CAGACAGATACTCCCGAACCTT *TFE3*-F: TGCGTCAGCAGCTTATGAGG *TFE3*-R: AGACACGCCAATCACAGAGAT

### Immunohistochemistry

Adult fly brains were dissected and fixed in 4% formaldehyde for 40 minutes, then washed with PBT (PBS + 0.1% TX-100) for 3 times and dissected further to remove additional debris in PBS solution. Clean and fixed brains were blocked in PBT solution with 5% Normal Donkey Serum (NDS) and subsequently stained with primary antibody at 4C overnight and then secondary antibody at room temperature for 2 hours. Primary antibody used: rat anti-DDC (1:300, Jay Hirsh). All secondary antibodies used were from Jackson ImmunoResearch (West Grove, PA, USA): donkey anti-rat Cy3 (1:1000, Cat. #712-165-153), donkey anti-mouse Cy5 (1:500, Cat. #715-175-150). All samples were mounted in Vectashield mounting media (Cat. #H-1500-10, Vector Laboratories, Burlingame, CA, USA).

### LysoTracker staining

Live adult fly brains were dissected in PBS, and immediately incubated in LysoTracker Red DND-99 (1:1000, Cat. # 40739ES50, Yeasen, Shanghai, China) solution with PBS (pre-warmed at 37°C) for 5 minutes at 25°C. Images were taken by the Nikon A1 microscope. Live IMG cells were seeded on 8-well chamber slides overnight, and incubated in LysoTracker Green DND-26 (1:10000, Cat. #L7526, Life technologies) or LysoTracker Red DND-99 with anti-fluorescence quenching DMEM (Cat. #A1896701, Gibco). Images were taken by Nikon Spinning Disk at 37°C in 5% CO2.

### Lysosomal integrity analysis

To analyze lysosomal integrity, we used a real-time imaging protocol of cells stained with acridine orange (AO, Cat. #ab270791, Abcam). IMG cells were incubated with AO (2 μg/ml) for 15 min at 37°C and rinsed with HBSS with 3% FBS. Cells from random areas were visualized and exposed to blue light for 20 s. Subsequently, images were captured for 10 min at 20-s intervals under the Nikon A1 microscope. By measuring the red color intensities, changes in lysosomal pH gradient were quantified.

### Cathepsin B detection with Magic Red

Live IMG cells were seeded on 8-well chamber slides overnight, and stained with 1X Magic Red Cathepsin B dye for 5 min, according to the manufacturer’s instruction (Magic Red^®^ Cathepsin B assay kits, Cat. #ICT937, BioRad). The red fluorescent intensities representing cathepsin B activity in the cells were monitored under the Nikon A1 microscope.

### Annexin V Staining

After *LRRK2* siRNA treatment, IMG cells were seeded on 8-well chamber slides overnight, and stained with 5 μl Annexin V-mCherry in 195μl Annexin V-mCherry Binding Buffer (Annexin V-mCherry Apoptosis Detection Kit, Cat. # C1069S, Beyotime, Shanghai, China). After incubation for 20 min in the dark at 37 °C in 5% CO2, the cells were captured under the Nikon A1 microscope.

### TUNEL Staining

IMG cells were seeded on glass coverslips in 12-well plates the day before fixation with cold methanol (Cat. #100141190, Sinopharm, Beijing, China) for 15 minutes, followed by washing with PBS. Adult fly brains were fixed and washed with similar protocol. Samples were then stained with Terminal Deoxynucleotidyl Transferase to detect cellular apoptosis according to the manufacturer’s protocol (One Step TUNEL Apoptosis Assay Kit, Cat. # C1089, Beyotime, Shanghai, China). The TUNEL-positive cells were observed by the Nikon C2 microscope.

### *In-vivo* time lapse analysis

Live adult fly brains were dissected and kept in saline solution for tracing the GFP puncta in a single focal plane using LSM 710 NLO (Zeiss, Germany) microscope with a 20X water objective. The trafficking speed was analyzed by Imaris using Brownian motion tracking mode.

### Transmission electron microscopy

Adult fly brains were fixed, embedded, stained, and dehydrated according to previous protocols(Li et al., 2013). Ultrathin sections (70 nm) of each brain were cut with a Diatomediamond knife on a Leica EM UC7 ultra-microtome (Leica Microsystems, Germany) and collected on copper grids. Images were acquired with Talos L120C transmission electron microscope (Thermo Fisher Scientific, Waltham, MA, UAS) operating at an acceleration voltage of 120 kV

### Behavior analysis

The rapid iterative negative geotaxis (RING) assay was used to analyze climbing ability as described previously (Cao et al., 2017; Gargano et al., 2005; Song et al., 2017). Briefly, flies of different ages (3-, 10-, or 20-day-old) were collected and placed in vials with fly food (no yeast) for 1 day before transferring to the cylinders for experiments. A total of 100 flies were analyzed in ten cylinders (inner diameter: 20 mm; height: 200 mm) placed side by side, and each contains ten unisex flies. An initial mechanical shock was applied for six times so that all flies were tapped down to the bottom of the cylinder and synchronized. Climbing distances for each fly were measured and averaged by RflyDetection software, which allows automatic detection of fly position within the cylinder using video images captured every 6 s (Sony digital camera, HDR-CX220E). At least three independent experiments were performed. Data were shown as mean ± SEM.

### Confocal microscopy and statistical analysis

Images of adult fly brains and IMG cells were acquired by scanning a serial Z-stack of average 10-15 sections, each of 0.6-1 μm thickness, using Nikon A1, C2 (Tokyo, Japan) with the 60X oil objective. The whole brain was positioned so that they can be scanned anteriorly to posteriorly (top to bottom). For statistical analyses of number and size, original and unmodified images were imported into ImageJ (National Institutes of Health), and the intensity threshold for the relevant channel was set so that maximum number of dots were selected without miscounting two adjacent dots into one. For colocalization measurements, primary images were imported and analyzed automatically using the colocalization plugin with the intensity correlation tool, showing as Pearson’s correlation coefficient (R). All data were imported into GraphPad Prism 8 and shown in scatter plots or column bar graphs. For calculating the statistical significance, two-tailed unpaired t-test or ordinary one-way ANOVA followed by Tukey’s multiple comparisons test was used. P value less than 0.05 is considered significant. ns: no significance, p≥ 0.05; *: p<0.05; **: p<0.01; ***: p<0.001; ****: p<0.0001.

## Figure Legends

**Figure S1.**
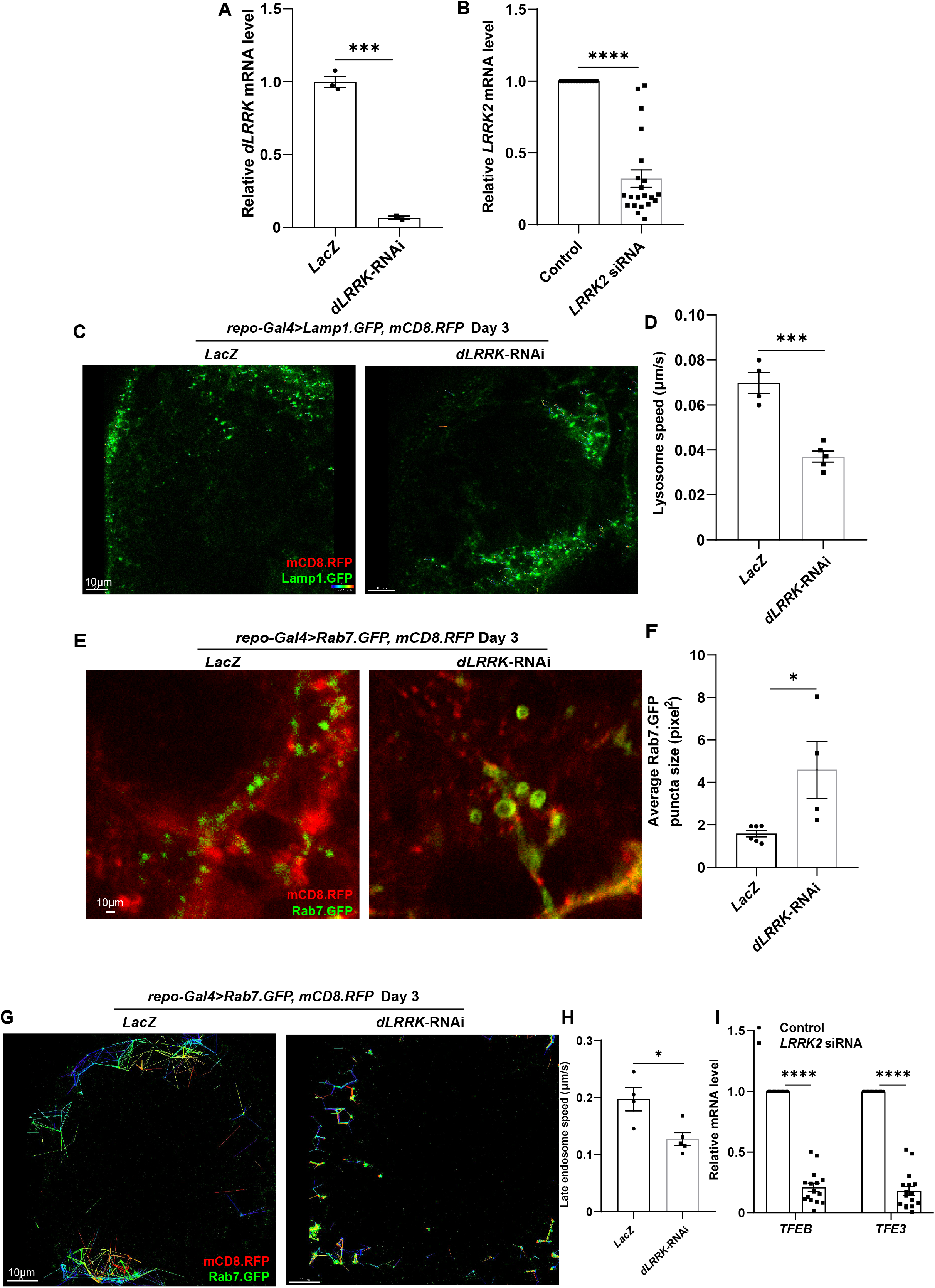
LRRK2/dLRRK mediates the glial lysosomal pathway. (**A** and **B**) Relative mRNA levels assessed by qRT-PCR upon glial LRRK2/dLRRK depletion in flies or IMG cells. Note that both expression of the *dLRRK*-RNAi or the treatment of *LRRK2* siRNA causes a dramatic decrease in the *LRRK2/dLRRK* mRNA level. (**C** and **D**) *In-vivo* time lapse analysis of glial lysosome mobility in adult fly brains with reduced *dLRRK* expression. Glial lysosomes (green) are labeled with *repo-GAL4* driven *UAS-Lamp1.GFP* while glial processes are marked with *UAS-mCD8.RFP* (*repo-Gal4*>*UAS-Lamp1.GFP*, *UAS-mCD8.RFP*). Note that the lysosome speed tracked with Imaris decreases in the absence of glial dLRRK. (**E** and **F**) Representative images (E) and quantifications (F) of Rab7.GFP-positive late endosomes in adult fly glia with the indicated genotypes. Note that glial late endosomes are also enlarged when expressing *dLRRK*-RNAi. (**G** and **H**) *In-vivo* time lapse analysis of glial late endosome mobility in adult fly brains with reduced *dLRRK* expression. Glial late endosomes (green) are labeled with *repo-GAL4* driven *UAS-Rab7.GFP* while glial processes are marked with *UAS-mCD8.RFP* (*repo-Gal4*>*UAS-Lamp1.GFP*, *UAS-mCD8.RFP*). Note that the late endosome speed tracked with Imaris decreases in the absence of glial dLRRK. (**I**) Relative mRNA levels of *TFEB* and *TFE3* assessed by qRT-PCR were decreased upon microglial LRRK2 depletion. Scale bars and the quantified sample number n are indicated as in Figure 1. Data are shown as mean ± SEM. P-values of significance (indicated with asterisks, ns no significance, * p<0.05, ** p<0.01, and *** p<0.001) are calculated by two-tailed unpaired t-test or ordinary one-way ANOVA followed by Tukey’s multiple comparisons test.

## References

Aasly, J.O., Vilarino-Guell, C., Dachsel, J.C., Webber, P.J., West, A.B., Haugarvoll, K., Johansen, K.K., Toft, M., Nutt, J.G., Payami, H., et al. (2010). Novel pathogenic LRRK2 p.Asn1437His substitution in familial Parkinson’s disease. Mov Disord 25, 2156–2163.

Baptista, M.A., Dave, K.D., Frasier, M.A., Sherer, T.B., Greeley, M., Beck, M.J., Varsho, J.S., Parker, G.A., Moore, C., Churchill, M.J., et al. (2013). Loss of leucine-rich repeat kinase 2 (LRRK2) in rats leads to progressive abnormal phenotypes in peripheral organs. PLoS One 8, e80705.

Beilina, A., Bonet-Ponce, L., Kumaran, R., Kordich, J.J., Ishida, M., Mamais, A., Kaganovich, A., Saez-Atienzar, S., Gershlick, D.C., Roosen, D.A., et al. (2020). The Parkinson’s Disease Protein LRRK2 Interacts with the GARP Complex to Promote Retrograde Transport to the trans-Golgi Network. Cell Rep 31, 107614.

Budnik, V., and White, K. (1988). Catecholamine-containing neurons in Drosophila melanogaster: distribution and development. J Comp Neurol 268, 400–413.

Cao, W., Song, L., Cheng, J., Yi, N., Cai, L., Huang, F.D., and Ho, M. (2017). An Automated Rapid Iterative Negative Geotaxis Assay for Analyzing Adult Climbing Behavior in a Drosophila Model of Neurodegeneration. Journal of visualized experiments: JoVE.

Cogo, S., Manzoni, C., Lewis, P.A., and Greggio, E. (2020). Leucine-rich repeat kinase 2 and lysosomal dyshomeostasis in Parkinson disease. J Neurochem 152, 273–283.

Cuenca, L., Gil-Martinez, A.L., Cano-Fernandez, L., Sanchez-Rodrigo, C., Estrada, C., Fernandez-Villalba, E., and Herrero, M.T. (2018). Parkinson’s disease: a short story of 200 years. Histol Histopathol, 18073.

Deng, X., Dzamko, N., Prescott, A., Davies, P., Liu, Q., Yang, Q., Lee, J.D., Patricelli, M.P., Nomanbhoy, T.K., Alessi, D.R., et al. (2011). Characterization of a selective inhibitor of the Parkinson’s disease kinase LRRK2. Nat Chem Biol 7, 203–205.

Di Fonzo, A., Rohe, C.F., Ferreira, J., Chien, H.F., Vacca, L., Stocchi, F., Guedes, L., Fabrizio, E., Manfredi, M., Vanacore, N., et al. (2005). A frequent LRRK2 gene mutation associated with autosomal dominant Parkinson’s disease. Lancet 365, 412–415.

Di Maio, R., Hoffman, E.K., Rocha, E.M., Keeney, M.T., Sanders, L.H., De Miranda, B.R., Zharikov, A., Van Laar, A., Stepan, A.F., Lanz, T.A., et al. (2018). LRRK2 activation in idiopathic Parkinson’s disease. Sci Transl Med 10.

Erb, M.L., and Moore, D.J. (2020). LRRK2 and the Endolysosomal System in Parkinson’s Disease. J Parkinsons Dis 10, 1271–1291.

Feany, M.B., and Bender, W.W. (2000). A Drosophila model of Parkinson’s disease. Nature 404, 394–398.

Gargano, J.W., Martin, I., Bhandari, P., and Grotewiel, M.S. (2005). Rapid iterative negative geotaxis (RING): a new method for assessing age-related locomotor decline in Drosophila. Experimental gerontology 40, 386–395.

Giaime, E., Tong, Y., Wagner, L.K., Yuan, Y., Huang, G., and Shen, J. (2017). AgeDependent Dopaminergic Neurodegeneration and Impairment of the Autophagy-Lysosomal Pathway in LRRK-Deficient Mice. Neuron 96, 796–807 e796.

Greggio, E., Jain, S., Kingsbury, A., Bandopadhyay, R., Lewis, P., Kaganovich, A., van der Brug, M.P., Beilina, A., Blackinton, J., Thomas, K.J., et al. (2006). Kinase activity is required for the toxic effects of mutant LRRK2/dardarin. Neurobiol Dis 23, 329–341.

Hasan, N., Greenwood, T., and Glunde, K. (2008). Lysosome swelling precedes lysosomal membrane permeabilization in breast cancer cells. Cancer Research 68, LB-51–LB-51.

Haugarvoll, K., Rademakers, R., Kachergus, J.M., Nuytemans, K., Ross, O.A., Gibson, J.M., Tan, E.K., Gaig, C., Tolosa, E., Goldwurm, S., et al. (2008). Lrrk2 R1441C parkinsonism is clinically similar to sporadic Parkinson disease. Neurology 70, 1456–1460.

Hayes, M.T. (2019). Parkinson’s Disease and Parkinsonism. Am J Med 132, 802–807.

Henry, A.G., Aghamohammadzadeh, S., Samaroo, H., Chen, Y., Mou, K., Needle, E., and Hirst, W.D. (2015). Pathogenic LRRK2 mutations, through increased kinase activity, produce enlarged lysosomes with reduced degradative capacity and increase ATP13A2 expression. Hum Mol Genet 24, 6013–6028.

Herzig, M.C., Kolly, C., Persohn, E., Theil, D., Schweizer, T., Hafner, T., Stemmelen, C., Troxler, T.J., Schmid, P., Danner, S., et al. (2011). LRRK2 protein levels are determined by kinase function and are crucial for kidney and lung homeostasis in mice. Hum Mol Genet 20, 4209–4223.

Hinkle, K.M., Yue, M., Behrouz, B., Dachsel, J.C., Lincoln, S.J., Bowles, E.E., Beevers, J.E., Dugger, B., Winner, B., Prots, I., et al. (2012). LRRK2 knockout mice have an intact dopaminergic system but display alterations in exploratory and motor coordination behaviors. Mol Neurodegener 7, 25.

Kuwahara, T., Inoue, K., D’Agati, V.D., Fujimoto, T., Eguchi, T., Saha, S., Wolozin, B., Iwatsubo, T., and Abeliovich, A. (2016). LRRK2 and RAB7L1 coordinately regulate axonal morphology and lysosome integrity in diverse cellular contexts. Sci Rep 6, 29945.

Li, H., Li, Y., Lei, Z., Wang, K., and Guo, A. (2013). Transformation of odor selectivity from projection neurons to single mushroom body neurons mapped with dual-color calcium imaging. Proc Natl Acad Sci U S A 110, 12084–12089.

Lin, C.H., Tzen, K.Y, Yu, C.Y., Tai, C.H., Farrer, M.J., and Wu, R.M. (2008). LRRK2 mutation in familial Parkinson’s disease in a Taiwanese population: clinical, PET, and functional studies. J Biomed Sci 15, 661–667.

McCarthy, R.C., Lu, D.Y, Alkhateeb, A., Gardeck, A.M., Lee, C.H., and Wessling-Resnick, M. (2016). Characterization of a novel adult murine immortalized microglial cell line and its activation by amyloid-beta. J Neuroinflammation 13, 21.

Munoz, L., Kavanagh, M.E., Phoa, A.F., Heng, B., Dzamko, N., Chen, E.J., Doddareddy, M.R., Guillemin, G.J., and Kassiou, M. (2015). Optimisation of LRRK2 inhibitors and assessment of functional efficacy in cell-based models of neuroinflammation. Eur J Med Chem 95, 29–34.

Nassel, D.R., and Elekes, K. (1992). Aminergic neurons in the brain of blowflies and Drosophila: dopamine- and tyrosine hydroxylase-immunoreactive neurons and their relationship with putative histaminergic neurons. Cell Tissue Res 267, 147–167.

Paisan-Ruiz, C., Jain, S., Evans, E.W., Gilks, W.P., Simon, J., van der Brug, M., Lopez de Munain, A., Aparicio, S., Gil, A.M., Khan, N., et al. (2004). Cloning of the gene containing mutations that cause PARK8-linked Parkinson’s disease. Neuron 44, 595–600.

Rocha, E.M., De Miranda, B.R., Castro, S., Drolet, R., Hatcher, N.G., Yao, L., Smith, S.M., Keeney, M.T., Di Maio, R., Kofler, J., et al. (2020). LRRK2 inhibition prevents endolysosomal deficits seen in human Parkinson’s disease. Neurobiol Dis 134, 104626.

Rudenko, I.N., and Cookson, M.R. (2014). Heterogeneity of leucine-rich repeat kinase 2 mutations: genetics, mechanisms and therapeutic implications. Neurotherapeutics 11, 738–750.

Sheng, Z., Zhang, S., Bustos, D., Kleinheinz, T., Le Pichon, C.E., Dominguez, S.L., Solanoy, H.O., Drummond, J., Zhang, X., Ding, X., et al. (2012). Ser1292 autophosphorylation is an indicator of LRRK2 kinase activity and contributes to the cellular effects of PD mutations. Sci Transl Med 4, 164ra161.

Song, L., He, Y., Ou, J., Zhao, Y., Li, R., Cheng, J., Lin, C.H., and Ho, M.S. (2017). Auxilin Underlies Progressive Locomotor Deficits and Dopaminergic Neuron Loss in a Drosophila Model of Parkinson’s Disease. Cell Rep 18, 1132–1143.

Steger, M., Tonelli, F., Ito, G., Davies, P., Trost, M., Vetter, M., Wachter, S., Lorentzen, E., Duddy, G., Wilson, S., et al. (2016). Phosphoproteomics reveals that Parkinson’s disease kinase LRRK2 regulates a subset of Rab GTPases. Elife 5.

Sveinbjornsdottir, S. (2016). The clinical symptoms of Parkinson’s disease. J Neurochem 139 Suppl 1, 318–324.

Tong, Y., Giaime, E., Yamaguchi, H., Ichimura, T., Liu, Y., Si, H., Cai, H., Bonventre, J.V., and Shen, J. (2012). Loss of leucine-rich repeat kinase 2 causes age-dependent biphasic alterations of the autophagy pathway. Mol Neurodegener 7, 2.

Tong, Y., Yamaguchi, H., Giaime, E., Boyle, S., Kopan, R., Kelleher, R.J., 3rd, and Shen, J. (2010). Loss of leucine-rich repeat kinase 2 causes impairment of protein degradation pathways, accumulation of alpha-synuclein, and apoptotic cell death in aged mice. Proc Natl Acad Sci U S A 107, 9879–9884.

Wang, F., Gomez-Sintes, R., and Boya, P. (2018). Lysosomal membrane permeabilization and cell death. Traffic 19, 918–931.

West, A.B., Moore, D.J., Biskup, S., Bugayenko, A., Smith, W.W., Ross, C.A., Dawson, V.L., and Dawson, T.M. (2005). Parkinson’s disease-associated mutations in leucine-rich repeat kinase 2 augment kinase activity. Proc Natl Acad Sci U S A 102, 16842–16847.

West, A.B., Moore, D.J., Choi, C., Andrabi, S.A., Li, X., Dikeman, D., Biskup, S., Zhang, Z., Lim, K.L., Dawson, V.L., et al. (2007). Parkinson’s disease-associated mutations in LRRK2 link enhanced GTP-binding and kinase activities to neuronal toxicity. Hum Mol Genet 16, 223–232.

Zhang, Y., Chen, K., Sloan, S.A., Bennett, M.L., Scholze, A.R., O’Keeffe, S., Phatnani, H.P., Guarnieri, P., Caneda, C., Ruderisch, N., et al. (2014). An RNA-sequencing transcriptome and splicing database of glia, neurons, and vascular cells of the cerebral cortex. J Neurosci 34, 11929–11947.

Zhang, Y., Sloan, S.A., Clarke, L.E., Caneda, C., Plaza, C.A., Blumenthal, P.D., Vogel, H., Steinberg, G.K., Edwards, M.S., Li, G., et al. (2016). Purification and Characterization of Progenitor and Mature Human Astrocytes Reveals Transcriptional and Functional Differences with Mouse. Neuron 89, 37–53.

Zimprich, A., Biskup, S., Leitner, P., Lichtner, P., Farrer, M., Lincoln, S., Kachergus, J., Hulihan, M., Uitti, R.J., Calne, D.B., et al. (2004). Mutations in LRRK2 cause autosomal-dominant parkinsonism with pleomorphic pathology. Neuron 44, 601–607.

